# Stem-cell modeling of cerebellar dysfunction of Angelman syndrome

**DOI:** 10.1101/2025.08.07.669046

**Authors:** Carina Maranga, Adriana A. Vieira, João Camões dos Santos, Teresa P. Silva, Joana Gonçalves-Ribeiro, Karim Chebli, Miguel Casanova, Maud Borensztein, Laura Steenpass, Sandra H. Vaz, Tiago G. Fernandes, Simão T. da Rocha, Evguenia P. Bekman

## Abstract

Angelman syndrome (AS) is a severe neurodevelopmental disorder caused by the loss of function of the maternal allele of the neuron-specific imprinted gene *UBE3A*. Although AS presents with developmental delay, ataxia, epilepsy, and speech impairment, the distinct roles of various brain regions in its pathology remain unclear, hindering the design of effective therapeutic strategies. Of all brain regions, the nature and impact of dysfunction of the cerebellum in AS remain elusive. To start addressing this, we leveraged our optimized stem cell-based model of regionally patterned human cerebellar organoids (hCerOs) capable of producing major cerebellar neuronal subtypes, including Purkinje and granule cells, further matured in a two-dimensional (2D) culture system. We modeled AS using induced pluripotent stem cell (iPSC) lines carrying the common class II deletion and unaffected controls previously validated in our system. hCerOs from both backgrounds recapitulate lineage commitment and neuron-specific *UBE3A* imprinting. AS hCerOs consistently show reduced size, impaired neuroepithelial expansion, and diminished expression of glutamatergic and GABAergic progenitor markers. Transcriptomic profiling and functional assays assessing neuronal activity during the maturation stage uncovered delayed maturation, abnormal firing and increased neuronal excitability in AS cultures. These results position our hCerO model as a robust platform for future mechanistic studies and therapeutic screening aimed at targeting cerebellar dysfunction in AS.

## Introduction

Angelman syndrome (AS) is a severe neurodevelopmental disorder characterized by intellectual disability, ataxia, speech impairment, epilepsy, hyperactivity, and a distinct happy demeanor ((Maranga et al., 2020; Margolis et al., 2015). This disease results from the loss of function of the maternally inherited *UBE3A* gene, which encodes an E3 ubiquitin ligase essential for protein homeostasis (Kishino et al., 1997; Matsuura et al., 1997). *UBE3A* expression is governed by genomic imprinting, being restricted to the maternal allele in neurons, while the paternal allele is silenced by the antisense transcript UBE3A-ATS, part of a longer SNHG14 polycistronic transcript (Rougeulle et al., 1997; Runte et al., 2001). Among the various molecular causes of AS, the most common is maternal deletions of the chromosome 15q11–q13 region, including class II deletions (∼5 Mb, ∼12 genes), which are associated with the most severe clinical phenotype (Keute et al., 2021; Yang et al., 2021), likely due to the combined effect of *UBE3A* loss and haploinsufficiency of the co-deleted genes.

Most insights into AS pathophysiology derive from mouse models, especially maternal *Ube3a* knockouts (KOs) (Jiang et al., 1998; Miura et al., 2002), which recapitulate some features of AS but exhibit milder phenotypes than humans, especially regarding motor coordination (Sonzogni et al., 2018). Compared to the larger deletions typically observed in humans, available mouse models involve more limited gene deletions, which along with species-specific differences in brain structure and function, may explain these discrepancies (Jiang et al., 1998; Jiang et al., 2010; Maranga et al., 2020; Miura et al., 2002). Differences in brain structure and development are considerable for the cerebellum, an area of the brain with a major role in motor coordination. Indeed, the development of the human cerebellum is more protracted and results in a larger, more intricately folded structure, whereas the mouse cerebellum develops faster and is structurally simpler with fewer folia (Haldipur et al., 2022). In mice, Purkinje cell-specific deletions of *Ube3a* do not recapitulate locomotor deficits (Dindot et al., 2007), whereas the constitutive maternal *Ube3a* KO model shows decreased tonic inhibition in granule cells, linked to impaired GABA transporter 1 (GAT1) function and associated motor deficits (Egawa et al., 2012). In humans, magnetic resonance imaging (MRI) and postmortem findings showed cerebellar atrophy and Purkinje cell abnormalities, potentially contributing to motor symptoms such as ataxia and coordination deficits (Du et al., 2023; Harting et al., 2009; Jay et al., 1991).

Species-specific differences in cerebellar development and complexity, along with the scarcity of human data highlight the need for human-based experimental systems to study cerebellar contributions to AS. Advances in pluripotent stem cell (PSC) and organoid technologies now offer powerful platforms to model human neurodevelopment *in vitro* with unprecedented fidelity (Birtele et al., 2025; Miura et al., 2022; Qian et al., 2016). While forebrain organoids have been used to explore aspects of cerebral cortex development and function in AS (Estridge et al., 2025; Sen et al., 2020; Sun et al., 2019) the cerebellum has remained overlooked in these efforts.

Here, we apply, for the first time, a guided, regionalized human cerebellar organoid (hCerO) model (Silva et al., 2020a) to induced PSCs (iPSCs) derived from two AS individuals with class II deletions and unaffected controls. We characterize the timing of *UBE3A* silencing during cerebellar differentiation and uncover consistent structural, molecular, and functional impairments in AS hCerOs and cerebellar neurons. Our findings point toward a role for cerebellar dysfunction in AS and propose this platform as a valuable tool for investigating disease mechanisms and testing therapeutic strategies, ultimately advancing our understanding of AS.

## Results

### AS-derived hCerOs generate all major lineages of cerebellar progenitors and neurons

To model cerebellar dysfunction in AS, we generated hCerOs using patient-derived iPSC lines with class II deletions (Fig. 1A), ASD (Pólvora-Brandão et al., 2018) and AG1.0 (Chamberlain et al., 2010), as well as unaffected control cell lines, F002.1A.13 (Silva et al., 2020a) and iPSC6.2 (Burridge et al., 2011) (Table S1). The initial part consists of a three-dimensional (3D) culture system (Fig. 1B) spanning 35 days which involves several stages: neural induction (1-7 days), specification of cerebellar territory (7-14 days), and cerebellar patterning (14-35 days). This is followed by a dissociation step, where day 35 hCerOs are enzymatically separated into single cells and subsequently reseeded in a two-dimensional (2D) culture system to enhance the differentiation and specialization of distinct neuronal subtypes, as outlined in the published protocol (Silva et al., 2020b), with minor corrections (see Materials and Methods).

**Figure 1.**
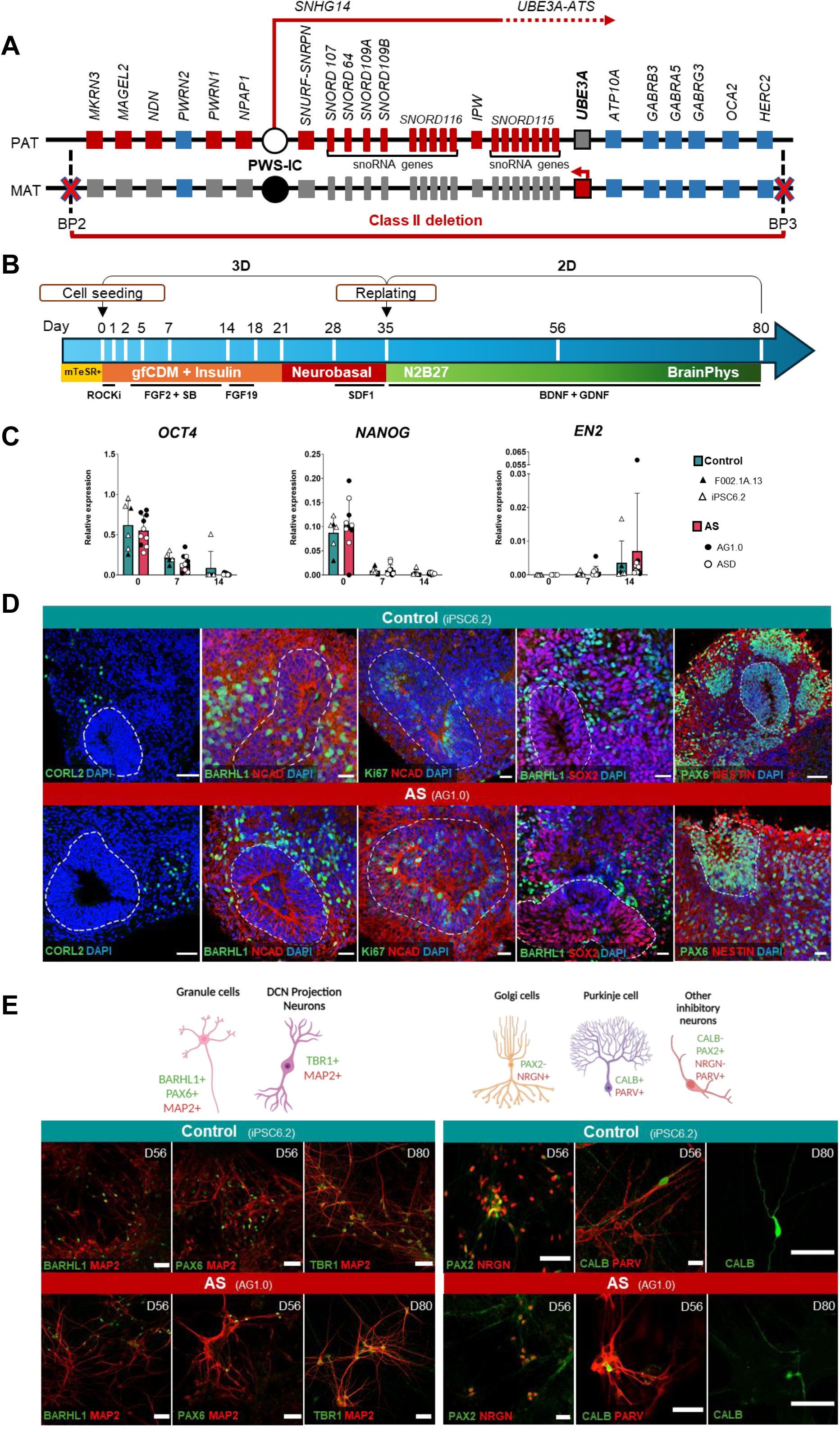
Correct patterning of cerebellar organoids from Angelman syndrome-derived iPSCs. **A)** Schematic representation of the 15q11-13 imprinted locus and respective class II deletion present in AG1.0 and ASD Angelman Syndrome (AS) iPSC lines. PAT: paternal chromosome; MAT: maternal chromosome; PWS-IC: Prader-Willi Syndrome-Imprinting Center; BP: breakpoint regions 2 (BP2) and 3 (BP3), leading to megadeletions; White circle: unmethylated region; Black circle: methylated region; Blue square: biallelically expressed gene; Red square: monoallelically expressed gene; Grey square: non-expressed gene. **B)** Schematic representation of the cerebellar differentiations. In the first phase, iPSCs were expanded in mTesR Plus, induced to aggregate in AggreWells^TM^ plates until day 7 and allowed to develop into 3D organoids. During differentiation two different culture media were used sequentially, growth factor-free chemically defined medium (gfCDM) and Neurobasal medium. On day 2, insulin, FGF2 and SB431542 were added. From day 14 to 21, FGF19 and on day 28 SDF1 was added. At day 35, organoids were dissociated and replated in a two-dimensional (2D) culture system in N2B27 medium into poly-ornithine and laminin coated wells. Starting from day 64, BrainPhys was progressively added to the cultures. Both N2B27 and BrainPhys media were supplemented with BDNF, GDNF and cAMP. **C)** RT-qPCR analysis of the represented genes in cerebellar organoids derived from F002.1A.13 and iPSC6.2 lines (unaffected controls) and AG1.0 and ASD iPSC lines (AS) at the indicated time-points. Data are presented as barplots representing the mean ± SEM of n ≥ 5 independent experiments per genotype; datapoints belonging to each cell line are indicated in the legend. Two-way ANOVA test followed by Sidak’s multiple comparison test was used for statistical analysis and no differences between AS and unaffected controls were found for the three genes. **D)** Representative immunofluorescence (IF) images of cerebellar organoids at day 35 for both control (iPSC6.2) and AS (AG1.0) organoids marked with CORL2 (in green) on the first panel; BARHL1 (in green) and NCAD (in red) in the second panel; Ki67 (in green) and NCAD (in red) in the third panel; BARHL1 (in green) and SOX2 (in red) in the fourth panel; PAX6 (in green) and NESTIN (in red) in the fifth panel; In all the panels, nuclei are counterstained with DAPI. White punctate lines outline neuroepithelial structures. Scale bars, 50 µm. **E)** On top, schematic representation of different type of cerebellar neurons [granule cells, deep cerebellar nuclei (DCN) projection neurons, Golgi cell, Purkinje cells and interneurons] and respective markers; In the bottom, representative IF images of glutamatergic (left) and GABAergic (right) markers in unaffected control (iPSC6.2) and AS-derived (AG1.0) cerebellar neurons at day 56 and 80. Scale bars, 50 μm.

The efficiency of neural induction was demonstrated by downregulation of pluripotency markers *OCT4* and *NANOG* by day 7 and upregulation of cerebellar patterning gene *EN2* by day 14 (Fig. 1C). By day 35, cerebellar organoids displayed a continuous cerebellar plate neuroepithelium with progenitors from two germinal zones: the ventricular zone (VZ), which gives rise to Purkinje cell progenitors and most cerebellar GABAergic interneurons, and the rhombic lip (RL), which generates granule cells, unipolar brush cells, and glutamatergic progenitors of deep cerebellar nuclei neurons (Muguruma et al., 2015). Representative immunofluorescence (IF) images of the day 35 hCerOs for AS (AG1.0) and control (iPSC6.2) demonstrate the presence of VZs exhibiting typical neuroepithelial organization (Fig. 1D, punctate outlines), with apical (internal) surface indicated by NCAD staining. Additionally, the presence of SOX2+, NCAD+, and Ki67+ GABAergic progenitors was observed (Fig. 1D). Differentiating CORL2+ Purkinje cell and GC precursors (BARHL1+, PAX6+) were located outside of the neuroepithelial structures, as expected. Importantly, hCerOs derived from AS patients did not display any signs of abnormal cerebellar patterning at this stage (Fig. 1D).

At later stages, replated AS and unaffected control cerebellar progenitors were able to generate all major types of cerebellar glutamatergic and GABAergic neurons (Fig. 1E), as evidenced by detection of granule cells co-expressing BARHL1/MAP2 and PAX6/MAP2, TBR1+/MAP2+ deep cerebellar nuclei projection neurons, CALB+/PARV+ Purkinje cells, PAX2-/NRGN+ Golgi cells, and other inhibitory interneurons (PAX2+/NRGN-, PARV+/CALB -) in both AS (AG1.0) and control (iPSC6.2) cultures at day 56 and day 80 of differentiation.

To sum up, we successfully differentiated both patient-derived and unaffected control iPSC lines into hCerOs, generating all major cerebellar neuronal lineages using our 80-day cerebellar differentiation protocol. These results enable us to move forward to the next critical step: evaluating whether *UBE3A* imprinting is established during this time window, an essential milestone for modeling AS.

### Cerebellar differentiation recapitulates imprinted UBE3A expression

During cerebellar differentiation, the paternal *UBE3A* allele is expected to become silenced, while expression from the maternal allele increases (Fig. 2A). In AS neurons lacking the maternal allele, *UBE3A* expression is therefore expected to be lost specifically in neurons (Fig. 2A). Accordingly, while we observed a continuous rise in *UBE3A* RNA levels during cerebellar differentiation of unaffected controls, these levels remained stagnant in AS iPSCs, as shown by RT-qPCR (Fig. 2B). Concomitantly, the neuron-specific *UBE3A-ATS* transcript, which is paternally expressed, shows a substantial increase in expression from day 56, marking the emergence of the first neurons in both AS and control cerebellar cultures. Accordingly, we observed reduced UBE3A IF signal in AS-derived cerebellar cultures in comparison to the control ones (Fig. 2C). As cerebellar cultures contain both non-neuronal (e.g., progenitor cells, astrocytes, etc) and neuronal cell populations, we performed double immunostaining for UBE3A and the neuronal marker MAP2 to confirm that, although UBE3A-positive cells persist in day 56 and day 80 AS cultures, they did not co-stain with MAP2 (Fig. 2D), In contrast in control cerebellar cultures, cells co-express UBE3A and MAP2 (Fig. 2D). Using Combined Bisulfite Restriction Analysis (COBRA), we confirmed that the methylation status of the Prader-Willi Syndrome-Imprinting Center (PWS-IC), essential for the imprinting regulation at the 15q11–q13 region, was maintained in both iPSCs and during differentiation. In unaffected controls, both the maternal methylated and paternal unmethylated alleles were present. In contrast, AS samples showed only the paternal unmethylated allele, indicating that the methylation marks at chromosome 15q11–q13 remained stable throughout differentiation, as expected (Fig. 2E). To further investigate when imprinted expression of *UBE3A* begins, we took advantage of the *UBE3A* mutation iPSC line, AS_Δ3 (Stanurova et al., 2016), that allows allele-specific distinction of *UBE3A* expression (Table S1). The AS_Δ3 iPSC line bears a 3 bp *de novo* in-frame deletion in exon 5 of the maternal *UBE3A* allele, that eliminates the G538 residue of UBE3A, providing a means to distinguish between the mRNAs produced from the two alleles. We employed pyrosequencing to determine the relative expression of the maternal and paternal alleles in day 0, 35, 56 and 80 cerebellar cultures derived from this cell line and established that the onset of paternal allele silencing occurs between days 35 and 56 (Fig. 2F). The allelic ratio on day 35 is approximately 50:50, but it shifts significantly by day 56 to 63:37 ± 2.71% (maternal:paternal) (Fig. 2F). This ratio remains largely unchanged on day 80 (61:39 ± 2.71%) (Fig. 2F), likely due to the increasing contribution of glial and other non-neuronal cells—particularly in AS cerebellar cultures—which corresponds with the observed reduction in *UBE3A-ATS* levels specifically in AS (Fig. 2B). Thus, we consider day 35 cultures to represent a largely pre-imprinting stage, and day 56 cultures a post-imprinting stage of *UBE3A* expression. Overall, these findings confirm that our cerebellar differentiation system accurately replicates the neuron-specific loss of UBE3A protein in AS cerebellar neurons.

**Figure 2.**
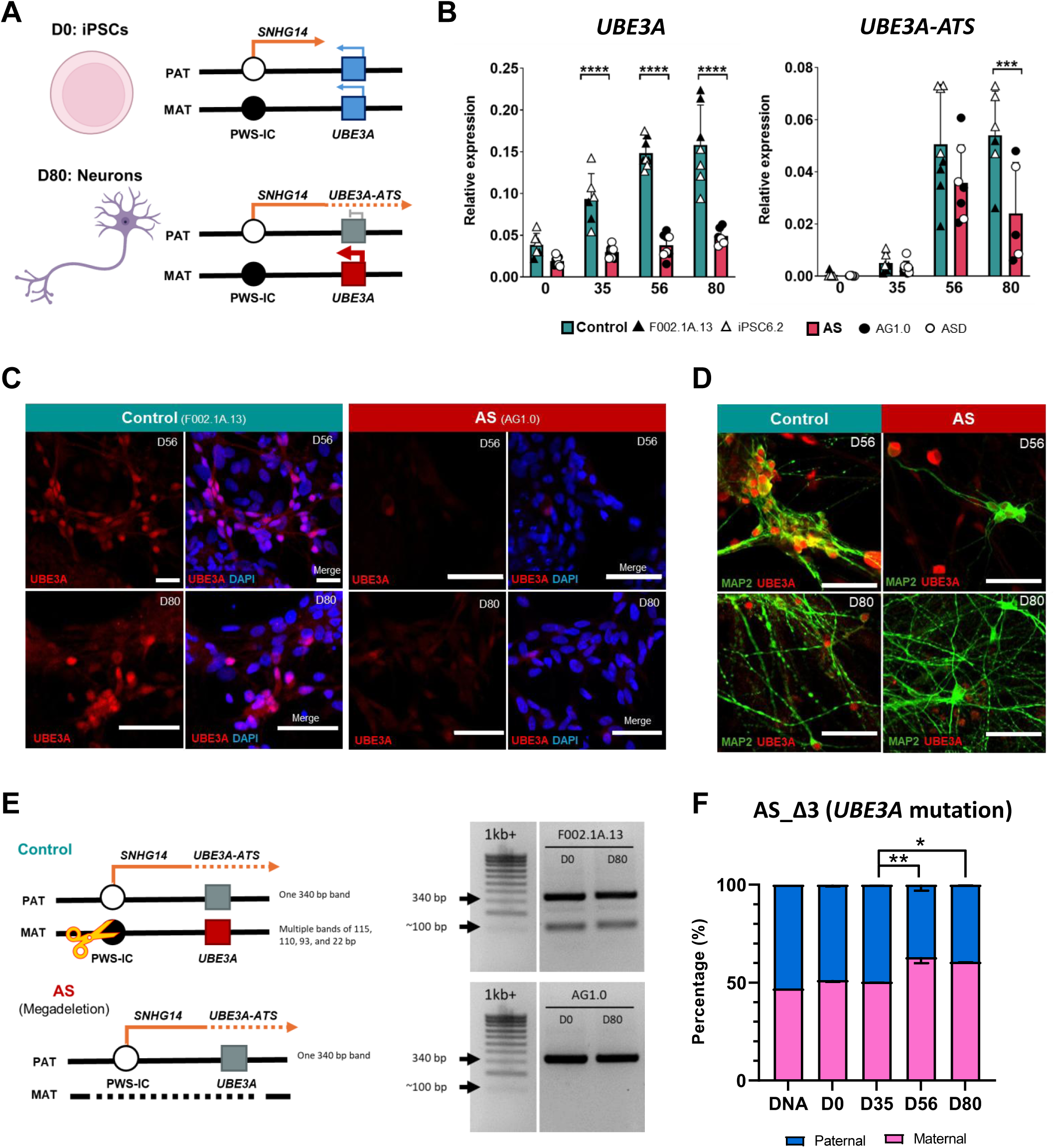
Imprinted expression of *UBE3A* during cerebellar differentiation. **A)** Schematic representation of allelic expression of *UBE3A* in iPSCs (day 0 of cerebellar differentiation) and neurons (day 80). PAT: paternal chromosome; MAT: maternal chromosome; PWS-IC: Prader-Willi Syndrome-Imprinting Center; White circle: un-methylated region; Black circle: methylated region; Blue square: biallelically expressed gene; Red square: monoallelically expressed gene; Grey square: silenced gene. Arrows indicate direction of expression and thickness of the arrow represent intensity of expression. Not drawn to scale. **B)** RT-qPCR analysis for *UBE3A* and *UBE3A-ATS* expression in unaffected control and AS-derived cultures, relative to *GAPDH* levels at the indicated time-points. Barplots show data as mean ± SEM for n ≥5 independent experiments per genotype; datapoints belonging to each cell line are indicated in the legend. Two-way ANOVA test followed by Sidak’s multiple comparisons test. Statistical significance: ***p<0.001; ****p<0.0001. **C)** Representative immunofluorescence (IF) images of UBE3A expression (in red) in cerebellar neurons derived from control (F002.1A.13) and AS (AG1.0) iPSCs at day 56 and 80. Nuclei counterstained with DAPI. Scale bars, 50 μm. **D)** Representative images of IF for MAP2 (in green) and UBE3A (in red) in control and AS iPSC-derived cerebellar differentiation at day 56 (F002.1A.13 and ASD) and 80 (F002.1A.13 & AG1.0). Scale bars, 50 μm. **E)** COmbined Bisulfite Restriction Analysis (COBRA) detecting methylation profile of the PWS-IC in the unaffected control (F002.1A.13) and in the AS (AG1.0) at D0 and D80 of cerebellar differentiation; on the left, the same scheme as in A displaying the expected size of the bands on the agarose gel for both the methylated and unmethylated PWS-IC; scissors indicate the the restriction enzyme used in the COBRA assay cleaves only the methylated PWS-IC; on the right, agarose gel shows the band profile for F002.1A.13 on top and AG1.0 in the bottom after COBRA. 1Kb+ column is the ladder with the indicated band sizes on the left. **F)** Allele-specific expression analysis by pyrosequencing of the *UBE3A* gene performed using the AS_Δ3 cell line at the indicated time points during cerebellar differentiation. The graph represents the % of maternal and paternal *UBE3A* alleles at different stages. Barplots shows the mean % ± SEM from n=3 independent experiments. Two-way ANOVA test followed by Tukey’s multiple comparisons test. Statistical significance: *p<0.05; **p<0.01.

### AS-derived cerebellar organoids exhibit smaller size, fewer neural rosettes and reduced expression of progenitor markers

After confirming that cerebellar differentiation with proper *UBE3*A imprinting can be achieved from both AS and control iPSCs, meeting essential criteria for modeling AS, we next examined potential phenotypic differences between AS and control hCerO and cerebellar neuronal cultures. The first parameter analysed was organoid size. Both AS iPSC lines consistently show smaller organoid size (Fig. 3A-B) by day 35 of differentiation. Analysis of the cryosections of hCerOs revealed the number of neural rosettes per organoid (Fig. 3C), as well as their size, is reduced in the AS condition, with statistically significant reduction of 5 out of 6 measured parameters, including apical and basal membrane length, as well as, total loop, loop tissue and ventricle areas (Fig. 3D-E). Quantifications of proliferation marker Ki67 (Fig. S1A) and progenitor marker SOX2 (Fig. S1B) within neuroepithelial structures, as well as the apoptosis marker cleaved CAS3 (Fig. S1C) showed no significant differences between the AS and control hCerOs. However, we noticed a slight increase in apoptosis in the AS condition by both IF (Fig. S2D) and flow cytometry analyses (Fig. S1E). Overall, these results indicate that while AS cerebellar organoids exhibit normal cerebellar patterning (Fig. 1D), they are smaller with fewer rosettes, a difference we could not attribute to significant changes in cell proliferation or apoptosis at day 35.

**Figure 3.**
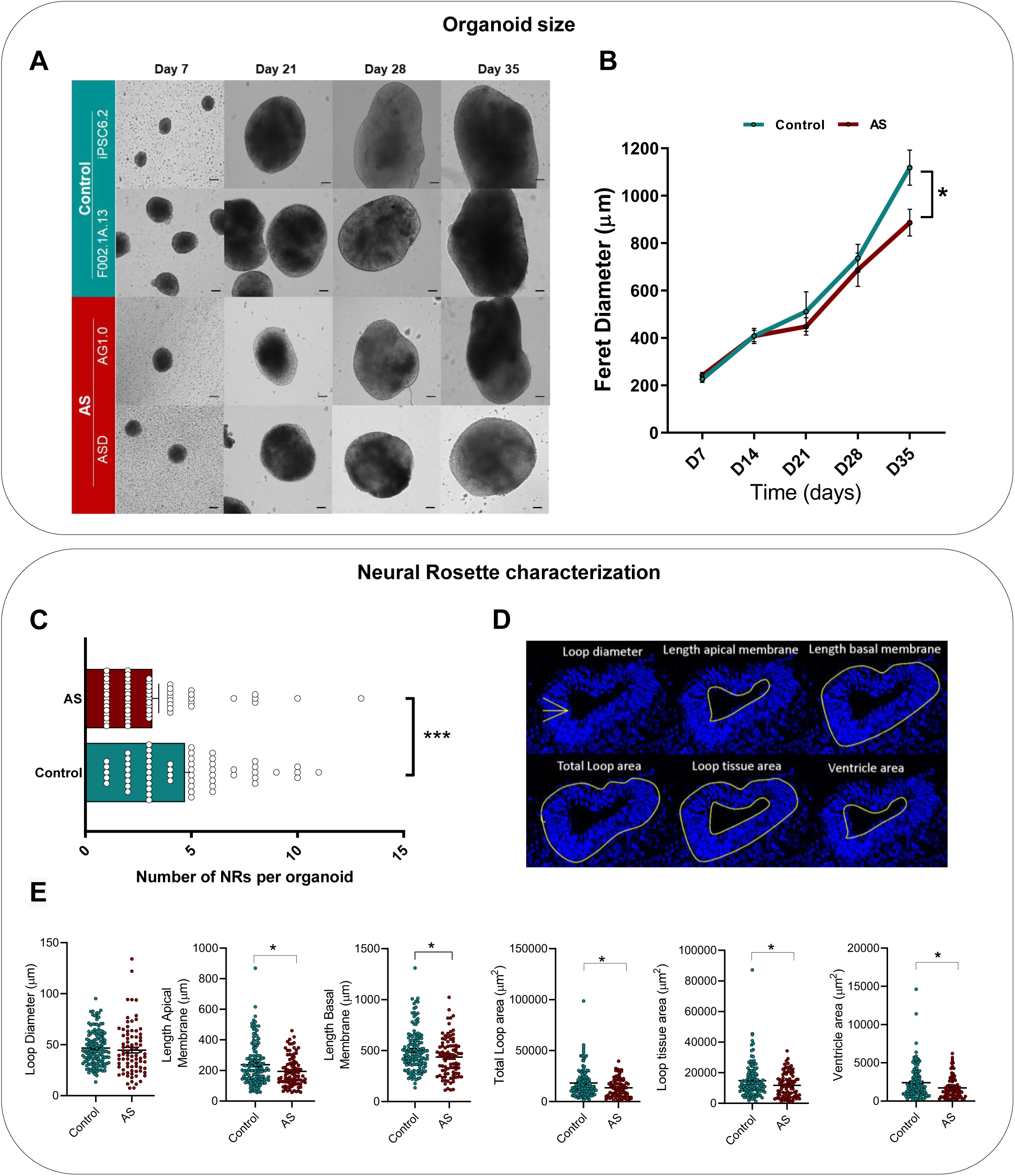
Reduced Organoid Size and Neural Rosette Formation in Angelman Syndrome Cerebellar Organoids. **A)** Representative bright-field microscopy images of control (iPSC6.2 & F002.1A.13) and AS (ASD & AG1.0) cerebellar organoids. Scale bars, 100 μm. **B)** Organoid size calculated through Feret diameter at indicated time points for control and AS cerebellar organoids. Line plot shows Mean size ± SEM of n ⩾ 7 organoids per genotype, including unaffected (F002.1A.13 and iPSC6.2) and AS (AG1.0 and ASD) hCerOs. Two-way ANOVA test followed by Sidak’s multiple comparisons test. Statistical significance: *p<0.05. **C)** Quantification of neural rosettes per organoid at day 35. Barplot represents the mean ± SEM of more than n ≥58 organoids per genotype, including unaffected (F002.1A.13 and iPSC6.2) and AS (AG1.0 and ASD) hCerOs, from at least 3 independent experiments. Mann-Whitney test. Statistical significance: ***p<0.001. **D)** Schematic illustrating quantification of morphological parameters: loop diameter, apical membrane length, total loop area, loop tissue area, and ventricle area. **E)** Morphological parameters as specified in D of neural rosettes evaluated at day 35 in control and AS-derived hCerOs. Barplot presents the mean ± SEM of ⩾80 neural rosettes per genotype, including unaffected (F002.1A.13 and iPSC6.2) and AS (AG1.0 and ASD) hCerOs from 8 independent experiments. Mann-Whitney test. Statistical significance: *p<0.05.

To explore this phenotype further, we next assessed in more detail the dynamics of progenitor cell specification in AS organoids. In accordance with the decreased size, AS-derived cerebellar organoids also exhibit an attenuated rise in the expression of pan-progenitor marker *NESTIN* during organoid specification/patterning (Fig. 4A). The GABAergic, VZ progenitor-associated genes *PAX2, OLIG2* and *KIRREL2,* as well as Purkinje cell precursor marker *CORL2,* all show lower mRNA levels in the AS-derived organoids (Fig. 4A), while among RL-associated progenitor markers (*PAX6, BARLH1* and *ATOH1*), only *PAX6* expression was significantly lower in AS condition (Fig. 4A). IF analyses for PAX2 and CORL2 show a similar trend with reduced number of positively stained cells for both GABAergic markers, similarly to the glutamatergic BARHL1 and PAX6, though only BARHL1 counts were significantly reduced in the AS condition (Figure 4B-C).

**Figure 4.**
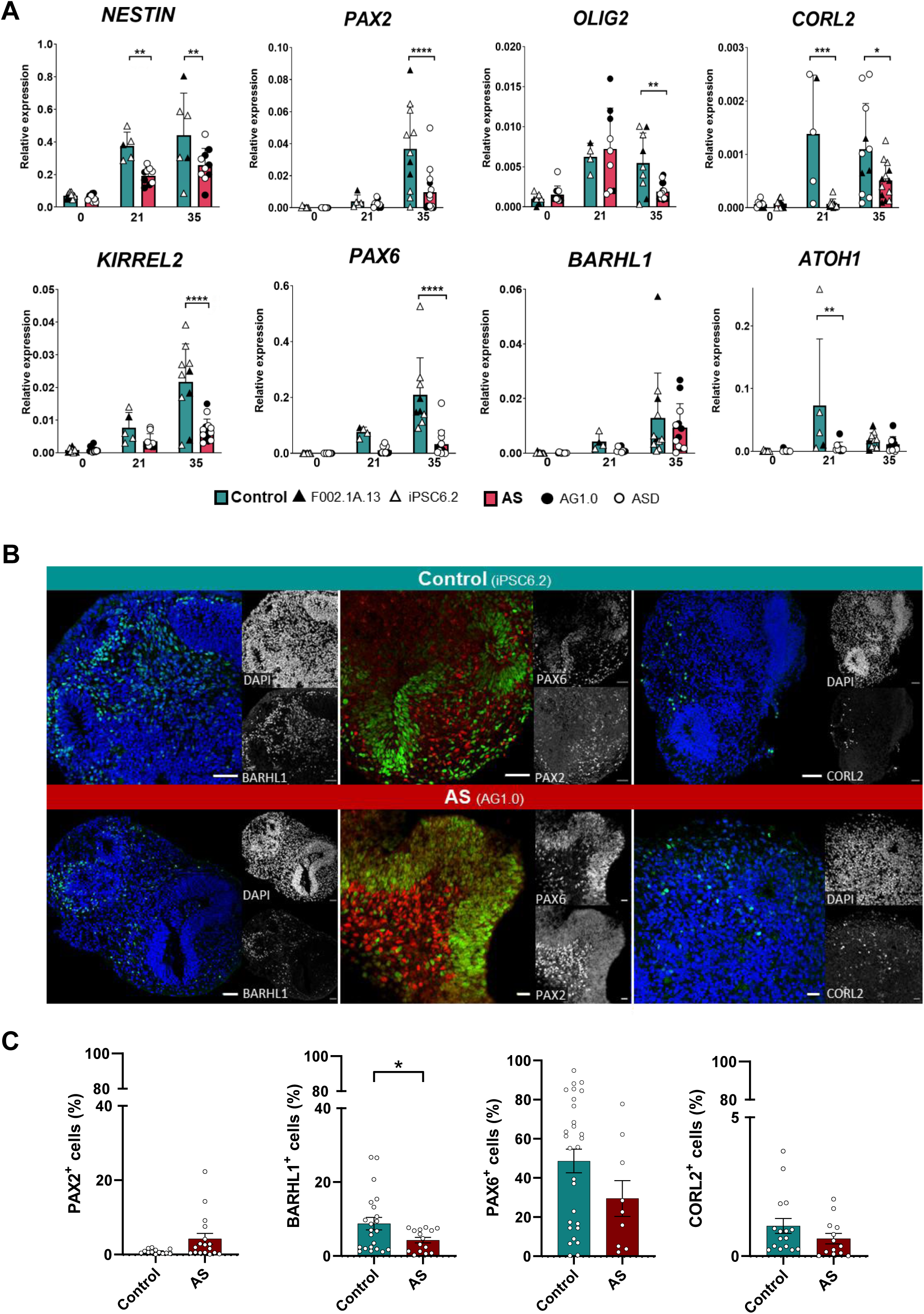
Reduced Expression of Cerebellar Progenitor Markers in Angelman syndrome-Cerebellar Organoids. **A)** RT-qPCR analysis of cerebellar progenitor markers in control and AS-derived cerebellar organoids at days 0, 21 and 35. Barplots represent the mean ± SEM of n ⩾ 4 independent experiments; datapoints belonging to each cell line are indicated in the legend below the barplots Two-way ANOVA test followed by Sidak’s multiple comparisons test. Statistical significance: *p<0.05; **p<0.01; ***p<0.001; ****p<0.0001. **B)** Representative immunofluorescence images of control (iPSC6.2) and AS-derived (AG 1.0) cerebellar organoids at day 35 for BARHL1 (in green) on the left, PAX6 (in green) and PAX2 (in red) in the middle, and CORL2 (in green) on the right; nuclei counterstained with DAPI. Scale bars, 50 μm. **C)** Fluorescent quantification for the indicated markers (PAX2, BARLH1, PAX6 and CORL2). Barplots show mean ± SEM quantified from ⩾ 9 of unaffected control and AS HCerOs from both unaffected control (F002.1A.13 & iPSC6.2) and AS (AG1.0 & ASD) cell lines. Statistically significant is represented as *p<0.05, unpaired two-tailed Student’s *t*-test or Mann-Whitney test.

In summary, smaller AS-derived organoids exhibit reduced expression of both GABAergic and glutamatergic progenitors, indicating a possible developmental delay and/or generation asynchrony in this cerebellar differentiation system. As this occurs prior to the first signs of *UBE3A* imprinting, it raises the possibility that haploinsufficiency of *UBE3A*, along with other genes deleted in class II deletions, may be relevant to the severity of this type of AS.

### Functional impairment of iPSC-derived cerebellar neurons in Angelman syndrome

We next investigated neuronal functionality in the replated monolayer cultures of AS and unaffected controls at days 56 and 80, after *UBE3A* imprinting has taken place and *UBE3A* silencing has occurred in AS cerebellar neurons (Fig. 2B,D,F). We started by performing single-cell calcium imaging (SCCI) to evaluate neuronal responses to stimuli by measuring changes in intracellular Ca²⁺ levels using Fura-2 dye (Gomes et al., 2020). To distinguish immature cells from neurons, intracellular Ca²⁺ responses were measured following stimulation with 100 μM histamine and 50 mM KCl, respectively (Agasse et al., 2008; Rodrigues et al., 2017) (Fig. 5A). On day 56, both AS and control cultures showed neuronal responses to KCl, however, the amplitude was markedly reduced in AS cultures (2–3-fold in controls vs. 1.5–2-fold in AS, Fig. 5Ai-ii). At this stage, control cultures responded well to histamine, whereas AS cultures showed a weaker response, potentially reflecting the reduced expression of progenitor markers observed at day 35 organoids. By day 80, KCl responses in AS neurons were nearly undetectable compared to the controls (Fig. 5Aiii-iv), while histamine-responsive cells increased, suggesting delayed maturation, selective progenitor survival and/or neuronal death. Importantly, non-responsive cells were always more prevalent in AS conditions, with the percentages of cells responding as neurons significantly decreasing by day 80 (Fig. 5B). Consistent with the SCCI analysis, immunostaining for TUJ1 and MAP2 confirmed a reduction in neuronal network density and a scarcity of mature PAX6⁺/MAP2⁺ granule cells as early as day 56 in AS cultures, along with a more pronounced increase in GFAP⁺ glial cells by day 80 (Fig. 5Ai-iv). Overall, these results indicate that AS cerebellar cultures exhibit a less developed and sparser neuronal network which declines with time in culture.

**Figure 5.**
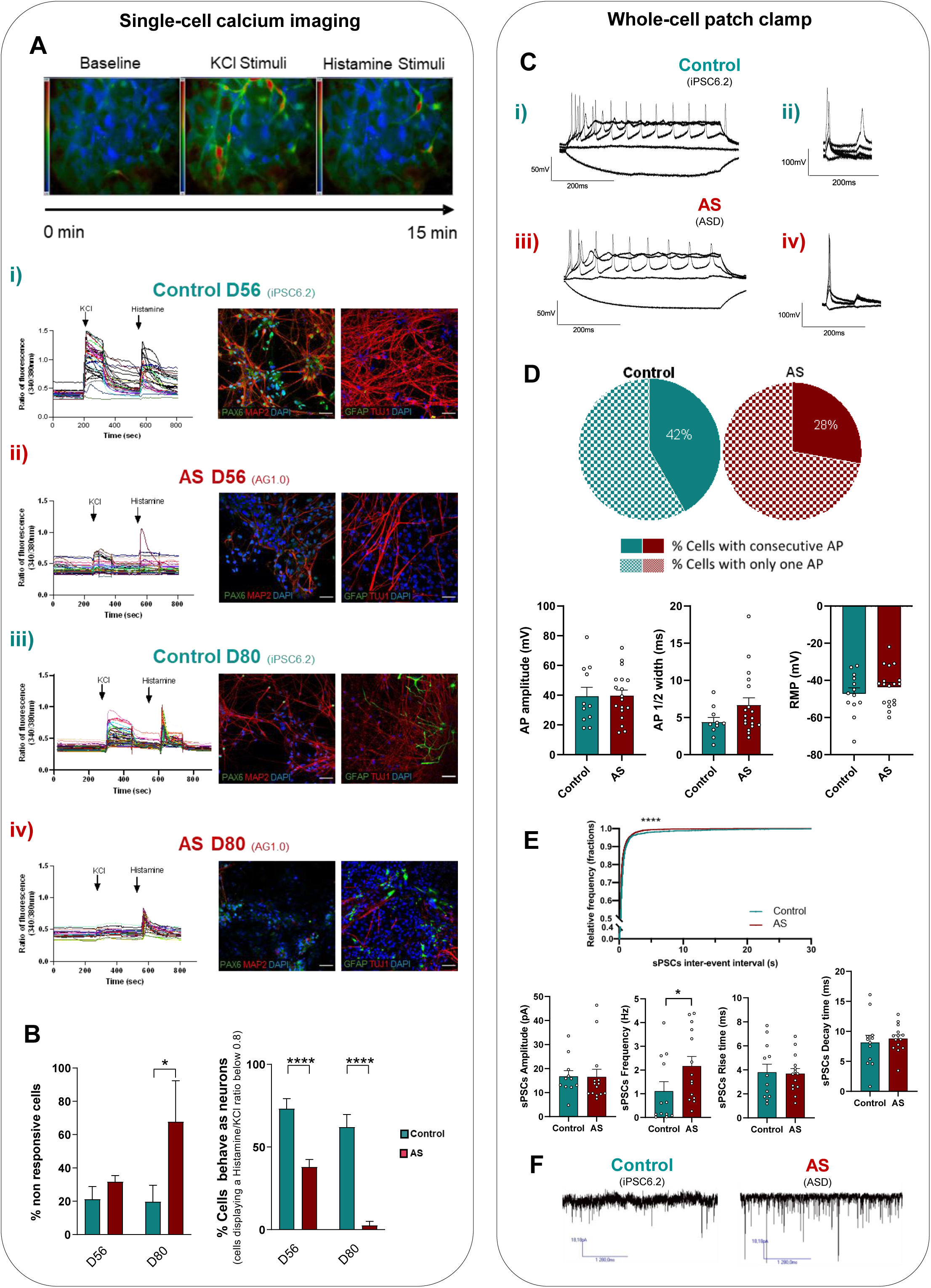
Functional Immaturity of AS Cerebellar Neurons Revealed by Electrophysiological Characterization of Cerebellar Culture. **A)** Single cell calcium imaging (SCCI): On top, representative fluorescent images of the applied protocol after potassium chloride (KCL) and histamine stimuli. In the bottom, representative fluorescence ratio profiles (left) of individual cells on days 56 and 80 for control (i, iii) and AS (ii, iv) are shown on the left. Data from both AS (AG1.0 & ASD) and unaffected control (F002.1A.13 & iPSC6.2) lines were combined. On the right, representative immunofluorescence images for PAX6 (in green), MAP2 (in red), TUJ1 (in red) and GFAP (in green) at days 56 and 80 in unaffected control (iPSC6.2) and AS (AG1.0) cerebellar cultures at day 56 and 80. **B)** Barplots represent the mean percentage of cells ± SEM that do not respond to changes in intracellular calcium levels when treated with KCl or histamine (left) and mean percentage ± SEM of responding cells that behave as mature neurons (cells displaying a Histamine/KCl ratio below 0.8; right). For day 56, 502 cells from 5 independent experiments were analysed for control and 539 cells from 6 independent experiments for AS. At day 80, 440 control cells from 5 independent experiments were evaluated and 51 AS cells from 2 independent experiments. For both day 56 and day 80, data from the two independent AS (AG1.0 & ASD) and control (iPSC6.2 & F002.1A.13) iPSC lines were pooled for analysis. Statistical significance is represented as *p<0.05, **** p<0.0001, two-way ANOVA test followed by Sidak’s multiple comparisons test. **C)** Top: Single-cell patch clamp recordings of single cells of control (iPSC6.2) and AS (ASD) cultures at day 80; i) Representative traces of firing responses evoked under current-clamp mode by injection of a 500 ms current pulse (−25 to +275 pA in 25-pA increments from an initial holding potential of −70 mV). Scale bars correspond to 50 mV and 200 ms. Right: Representative traces of firing responses to two independent current injections (10 ms) separated by 80 ms in control and AS-derived neurons. Scale bars correspond to 100 mV and 200 ms. **D)** Quantification of the electrical activity parameters obtained from patch-clamp recordings. AP amplitudes and half-peak width measured from the first AP depicted. Resting membrane potential (RMP). Circle graphs represent the percentage of cells with or without a consecutive AP after two independent current injections. For control were evaluated ⩾10 cells from 4 independent experiments. Data from both AS (AG1.0 & ASD) and unaffected control (F002.1A.13 & iPSC6.2) lines were combined. **E)** Quantification of the electrical parameters obtained from recordings of spontaneous postsynaptic currents. Frequency, amplitude, rise time, decay time of spontaneous events and sPSCs inter-event interval; data from 12 cells (4 independent experiments) and 14 cells (4 independent experiments) for control and AS, respectively; data from the two independent AS (AG1.0 & ASD) and control (iPSC6.2 & F002.1A.13) iPSC lines were pooled for analysis; Statistically significance is represented as *p<0.05 and ****p<0.0001, Unpaired two-tailed Student’s *t*-test or Mann Whitney test; error bars represent SEM. **F)** Representative recordings of spontaneous postsynaptic currents in control (iPSC6.2) and AS (ASD) cerebellar neurons. Scale bars correspond to 18.18 pA and 1280 ms.

We also assessed the electrophysiological properties of differentiated neurons using whole-cell patch-clamp recordings at day 80 of differentiation, when the consistent activity of mature neurons was detected according to the original protocol (Silva et al., 2020a). Both control and AS neurons presented typical firing action potentials (APs) upon a current injection (Fig. 5Ci, iii). However, after applying two independent current injections separated by 80 ms, 72% of AS neurons failed to respond to the second stimulus, in contrast to 58% of control ones (Fig. 5Cii, iv, and Fig. 5D). While no significant differences were observed in AP amplitude, half-peak width, and resting membrane potential (RMP) (Fig. 5D), AS condition exhibited higher frequency of spontaneous postsynaptic currents (sPSCs) (2.17 ± 3.99 Hz) compared with the control (1.11 ± 3.99 Hz) as shown in representative traces (Fig. 5E-F). This difference is not due to the amplitude, rise time, or decay time of sPSCs, but rather to their increased frequency, which results from shorter intervals between events (Fig. 5E-F). These findings indicate that cerebellar neurons in both conditions developed mature membrane properties and functional synapses by day 80, with the elevated spiking activity in AS-derived neurons aligning with the excitatory phenotype characteristic of the syndrome (Wallace et al., 2012). To complete functional characterization, dendritic spine analysis was performed, detecting overall reduced spine number per dendrite (Fig. S3A) with no major differences in individual spine density maturation (Fig. S3B-C). Overall, whole-cell patch-clamp recordings indicate an immature neuronal phenotype and an excitatory profile in AS cerebellar neurons.

### Transcriptomic profiling of cerebellar cultures at day 56 reflects neuronal immaturity of AS cerebellar neurons

To consolidate our functional analysis on AS neuronal cultures with molecular data, we performed transcriptomic profiling of AS and unaffected control cerebellar cultures on day 56, when AS cultures contain the highest number of functional neuronal cells (Fig. 5A-B). Bulk RNA-seq was performed in three replicates each for both control (2 F002.1A.13 & 1 iPSC6.2) and AS (2 AG1.0 & 1 ASD). Principal component analysis (PCA) showed variability between the samples with PC1 accounting for 45.4% of the variation, primarily distinguishing genotypes, and PC2 explaining 25.8%, separating lines within each genotype (Fig. 6A). In accordance with our RT-qPCR results (Fig. 2B), RNA-seq data show a statistically significant decrease in *UBE3A* expression and a tendency for decreased expression of *SNHG14*, the large polycistronic unit containing *UBE3A-ATS*, in AS cultures (Fig. S3A-B; Table S2). Differential gene expression analysis (DEA) (|log2foldchange| >1 and padj >0.05) detected 97 upregulated and 90 downregulated genes in AS compared to control cultures (Fig. 6B; Table S2). Amongst the statistically significant downregulated genes in AS cultures, we found *UBE3A*, zinc finger proteins (*e.g.*, *ZNF736*, *ZNF835* and *ZNF229*), and several non-coding genes. Several protein-coding genes, including *RGPD1* and *RGPD2* were found amongst the top upregulated genes in AS cultures. The relevance of these genes in the AS pathology remains to be investigated.

**Figure 6.**
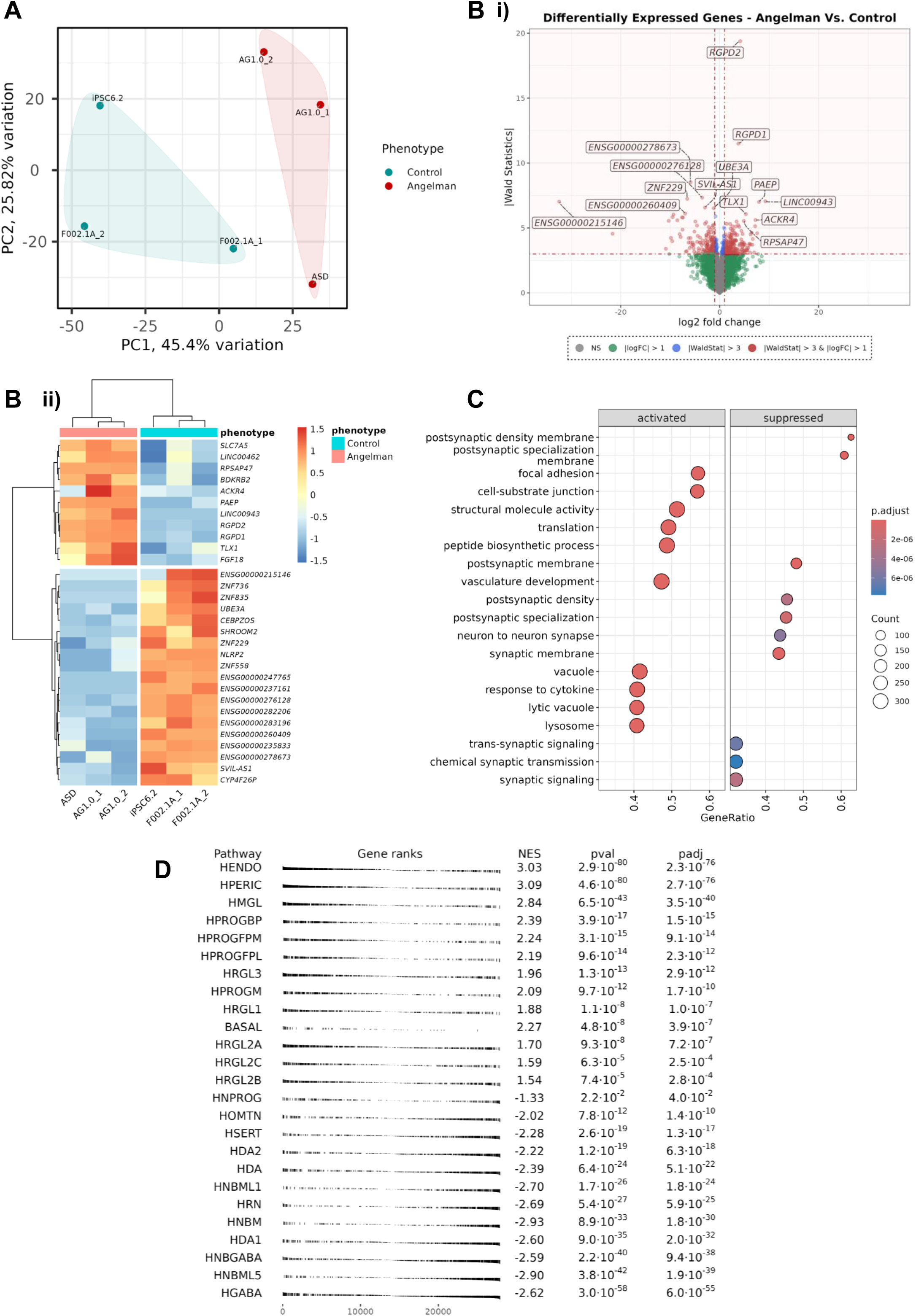
Transcriptomic Profiling Reveals Functional Immaturity in AS Cerebellar Neurons. **A)** Principal Component Analysis (PCA) of bulk RNA-seq data from day 56 cerebellar cultures derived from unaffected control (n=3; 2x F002.1A.13 and 1x iPSC6.2) and AS (n=3; 2x AG 1.0 and 1x ASD) lines. **B) i)** Volcano plot showing differentially expressed genes between AS-derived and control-derived cerebellar organoides at day 56 of differentiation. Genes with |log2 fold change| > 1 and Wald statistic > 3 are highlighted.**ii)** Heatmap showing the top 30 most differentially expressed genes in AS vs control cerebellar cultures. Gene expression is scaled across rows, and samples are annotated by phenotype. **C)** Dot plot representing a Gene Ontology (GO) enrichment analysis of differentially expressed genes, showing the top biological processes enriched among activated and suppressed gene sets in AS cultures. Dot size represents the number of genes associated with each term; color scale corresponds to adjusted p-values. **D)** Gene Set Enrichment Analysis (GSEA) using curated single-cell-derived neurodevelopmental transcriptional programs (La Manno et al., 2016). Enrichment was assessed in AS versus control day 56 cerebellar cultures. NES: Normalized Enrichment Score; pval: nominal p-value; padj: Benjamini-Hochberg adjusted p-value. Pathway acronyms correspond to specific neurotypes: HENDO (hindbrain endothelial cells), HPERIC (hindbrain pericytes), HMGL (hindbrain microglia), HPROGGBP, HPROGFPM, and HPROGFPL (hindbrain progenitor subtypes), HPROGM (midline progenitors), HRGL1,3 and HRGL2A–B (radial glia-like cells), BASAL (basal progenitors), HNPROG (hindbrain neuronal progenitors), HOMTN (hindbrain oligodendrocyte/mesodermal transitional neurons), HSERT (serotonergic neurons), HDA, HDA1, and HDA2 (hindbrain dopaminergic neurons), HNBML1 and HNBML5 (hindbrain neuronal branch-like neurons), HRN (hindbrain reticular neurons), HNBGABA (hindbrain GABAergic neurons), HNBM (hindbrain branch motor neurons), and HGABA (general GABAergic neurons).

Gene ontology (GO) enrichment analysis showed that upregulated gene pathways in AS cultures were largely unrelated, involving the distinct processes such as adhesion, translation, and RNA processing. In contrast, pathways associated with synaptic function and structure were consistently downregulated in AS cerebellar cultures (Fig. 6C). These findings are consistent with the functional immaturity observed in AS cerebellar neurons through SCCI and dendritic spine morphology analyses (Fig. 5A-B; Fig. S2). To investigate cell type-specific transcriptional alterations in AS, we leveraged a curated single-cell transcriptomic reference atlas from the developing human brain (La Manno et al., 2016). Using gene set enrichment analysis (GSEA), we assessed whether transcriptional signatures of distinct neuronal populations were differentially enriched in day 56 cerebellar cultures derived from AS versus unaffected controls (Fig. 6D). We observed, in AS cultures, a significant enrichment of gene expression programs associated with radial glia-like cells, neural progenitor populations, as well as non-neuronal types including pericytes and endothelial cells. In contrast, signatures characteristic of mature neuronal populations, were significantly depleted in AS samples. Overall, AS cultures display clear molecular signatures of neuronal immaturity, in line with our functional findings. This highlights the capacity of our cerebellar differentiation system to capture disease-relevant phenotypes specifically associated with the AS genotype.

## Discussion

Our study successfully generated hCerOs and cerebellar neuronal cell types from patient-derived AS iPSC lines with class II deletions, the most frequent cause of AS. These hCerOs exhibit normal cerebellar patterning at early stages and are able to recapitulate *UBE3A* imprinting once neurons emerge in culture. However, AS cerebellar organoids displayed reduced size, fewer neural rosettes, and lower expression of progenitor markers, indicating an early developmental delay, occurring before *UBE3A* imprinting takes place. At later stages, when UBE3A protein is no longer expressed in neurons, functional impairments reflecting an immature and hyperexcitable phenotype are detected in AS cerebellar neurons. These findings highlight the potential of patient-derived hCerOs as a relevant model for studying AS-specific cerebellar dysfunction and associated developmental abnormalities.

A key strength of our model is its ability to replicate the silencing of the paternal *UBE3A* allele in cerebellar neurons, a crucial feature for studying AS. Nonetheless, residual expression of paternal *UBE3A* persists in AS cerebellar neurons, with no significant reduction between day 56 and day 80 of differentiation (Fig. 2B,F). This sustained expression may reflect the overall immaturity of AS cerebellar cultures, the presence of non-neuronal cells, and a decline in the proportion of functionally active neurons by day 80 (Fig. 5A–B; Fig. 6C–D). Despite this, UBE3A protein levels remained consistently lower in AS cultures compared to controls (Fig. 2B), consistent with levels observed in other stem cell–based differentiation models (Stanurova et al., 2016; Sen et al., 2020). Together, these findings support the use of patient-derived hCerOs as a physiologically and disease-relevant model for investigating AS pathogenesis.

AS hCerOs displayed normal cerebellar patterning at day 35 (Fig. 1D), but were significantly smaller, with fewer neural rosettes per organoid (Fig. 3A–E) and reduced expression of progenitor markers (Fig. 4A–C). Notably, these phenotypes emerge before *UBE3A* imprinting becomes evident (Fig. 2F), suggesting that *UBE3A* haploinsufficiency, rather than complete loss, along with the hemizygous deletion of other genes in the class II region, may drive these early AS-specific traits. Although human data on embryonic cerebellar development in AS are limited, AS mouse models exhibit a smaller cerebellum at birth that normalizes by weaning (Jiang et al., 1998), consistent with delayed cerebellar development. Supporting this, a recent MRI study of 73 individuals with AS reported globally reduced grey matter volume, including significant reductions in the subcortical nuclei, cortex, and cerebellum (Du et al., 2023). Interestingly, while AS hCerOs exhibit early size reductions, such phenotypes were not reported in cortical organoids derived from homozygous *UBE3A* KO hESCs (Sun et al., 2019); however, these differences should be interpreted with caution given key distinctions between the models - cell source (hiPSC vs. hESC), genetic alteration (class II deletion vs. homozygous *UBE3A* KO), and brain region specificity (cerebellar vs. cortical differentiation protocols). Curiously, a recent study using a similar homozygous *UBE3A* knockout stem cell model reported contrasting findings in unguided cerebral organoids, which self-organize without regional patterning cues and contain mixed brain region identities (Estridge et al., 2025). They observed a reduction in organoid size and identified an impact of UBE3A loss on progenitor populations, as confirmed by single-cell RNA sequencing, findings that are consistent with our observations in AS hCerOs. Also in accordance with these and our results, Simchi et al. (Simchi et al., 2023) demonstrated that AS mouse-derived neural progenitors exhibit mitochondrial dysfunction, elevated ROS, and increased apoptosis. These phenotypes in neural progenitors arise prior to *UBE3A* imprinting and point to a dosage-dependent role of UBE3A in mitochondrial and oxidative stress pathways. Our findings of reduced organoid size and delayed development are consistent with these progenitor-stage deficits and suggest that *UBE3A* haploinsufficiency contributes to early pathogenic mechanisms in AS, potentially exacerbated by loss of neighboring genes in the deletion region. In sum, the reduction in progenitor populations observed in hCerOs is consistent with reduced grey matter volume in children with AS (Du et al., 2023), smaller organoid size and diminished progenitor proliferation in unguided cerebral organoids (Estridge et al., 2025) and altered progenitor dynamics reported by Simchi et al. (2023), underscoring the importance of investigating early-stage phenotypes in AS research.

By day 56, AS-derived cerebellar neurons displayed clear signs of immaturity, including reduced neuronal network complexity (Fig,5A), impaired synaptic function (Fig.5A-F), and a lower density of dendritic spines (Fig. S2A). These functional deficits are consistent with our transcriptomic data, which show enrichment for progenitor-associated signatures and a downregulation of mature neuron-specific genes (Fig. 6D). Gene ontology (GO) analysis further revealed a significant downregulation of pathways related to the post-synaptic density and post-synaptic membrane (Fig. 6C). Interestingly, although UBE3A is not localized to the post-synaptic compartment, it has been shown to accumulate in dendritic spines and the neuropil (Burette et al., 2018), suggesting a critical role in synaptic development and function. The observed synaptic abnormalities in AS cerebellar neurons thus support a role for UBE3A in the establishment and maintenance of synaptic integrity. Taken together, these functional deficits observed in iPSC-derived cerebellar neurons may be relevant to features of AS symptomatology, such as motor coordination impairments, ataxia, and cognitive deficits, although the extent of their contribution remains unclear. Another interesting phenotype, the increased synaptic excitability observed in AS-derived neurons (Fig. 5F) aligns with the hyperexcitability phenotype previously described in both AS animal models and patient-derived systems (Copping and Silverman, 2021; Fink et al., 2017; Judson et al., 2016; Sun et al., 2019; Wallace et al., 2017) and present in AS individuals, specifically the increase in EEG (electroencephalography) δ-power (Frohlich et al., 2019; Sidorov et al., 2017). Together, these findings highlight a potential contribution of cerebellar synaptic dysfunction to the pathophysiology of AS, underscoring the need for further studies into the cerebellum’s role in mediating disease symptoms.

While our model provides valuable insights into AS cerebellar development, several important limitations must be acknowledged. A major constraint is the need to dissociate 3D cerebellar organoids into 2D cultures to achieve long-term viability and neuronal maturation. This dissociation disrupts the native tissue architecture and may impair cell–cell interactions critical for proper cerebellar development. At the time this study was initiated, alternative protocols were not available; however, recent advances in organoid culture now allow intact hCerOs to be maintained for at least six months (Atamian et al., 2024; Atamian et al., 2025). Additionally, our *in vitro* system lacks the complex in vivo interactions between the cerebellum and other brain regions, and the short differentiation timeline limits its ability to model the protracted development seen in the human cerebellum. Incorporating strategies such as co-culturing with other brain regions (e.g., assembloids) or using microfluidic platforms may help better recapitulate physiological conditions. Finally, the robustness and generalizability of our findings must be validated in a broader panel of AS patient-derived stem cell lines, including isogenic controls and models representing different genetic subtypes of AS, to account for inter-individual variability and minimize potential off-target effects. These refinements will be essential to enhance the accuracy and translational relevance of this model for investigating AS pathophysiology.

Overall, our study highlights the potential of patient-derived hCerOs as a platform for studying AS-specific cerebellar dysfunctions. The identified molecular and functional alterations pave the way for future research into targeted therapeutic strategies. Further refinement of this model and validation of its findings with *in vivo* systems will be essential steps toward developing effective interventions for AS.

## Acknowledgments

This work was supported by Pedro Maria José de Mello Costa Duarte 2019 grant by Fundação Amélia de Mello, by the project*“Stem cell toolkit for modelling cerebellar dysfunction in Angelman syndrome”* granted in 2022 by Angelman Syndrome Alliance and by the CEREBEX Research grant LISBOA-01-0145-FEDER-029298 co-funded by FEDER (POR Lisboa 2020—Programa Operacional Regional de Lisboa PORTUGAL 2020) and Fundação para a Ciência e a Tecnologia (FCT).

Additional support by FCT and Ministério da Ciência, Tecnologia e Ensino Superior (MCTES) of Portugal came from the IC&DT projects, PTDC/BIA-MOL/29320/2017 and 2022.01532.PTDC, as well as projects UIDB/04565/2020 and UIDP/04565/2020 of the Research Unit Institute from Bioengineering and Biosciences – iBB and LA/P/0140/2020 of the Associate Laboratory Institute for Health and Bioeconomy – i4HB.

C.M. and J.C.d.S. were supported by the FCT PhD fellowships, PD/BD/135505/2018 and 2023.01932.BD, respectively.

Lastly, we acknowledge the Portuguese Angelman syndrome patient’s association Angel for their enthusiastic support of our work.

## Materials and Methods

### Ethics

Human iPSC lines were either purchased or previously generated by our team (Table S1). Donor consent was obtained, and ethical approval was granted by the Lisbon Academic Medical Center (Approval number: 535/12).

### Maintenance of iPSCs

The experiments were performed using karyotypically normal and pluripotent hiPSC lines derived from AS individuals with a class II deletion (ASD and AG1.0) and two unaffected and unrelated controls (F002.1A.13 and iPSC6.2) (Pólvora-Brandão et al. 2018; Chamberlain et al. 2010; Silva et al., 2020a; Burridge et al. 2011) (Table S1). iPSCs were cultured in six-well plates coated with Matrigel^TM^ (Corning, #354230) with mTeSR^TM^ Plus Medium (StemCell Technologies, #100-0276) at 37°C in a humidified 5% CO2 incubator. The medium was changed every other day. Cells were passed every three to four days using 0.5 mM EDTA dissociation buffer (Thermo Fisher Scientific, #15575-038). Cells underwent two to three passages prior to the initiation of the differentiation protocol.

### 3D culture of cerebellar organoids

Cerebellar differentiation was performed as described in (Silva et al. 2020) with minor modifications. Briefly, iPSCs were dissociated with Accutase (Sigma, #A6964) for 5 min at 37°C and re-aggregated in microwell plates (AggreWell™800, StemCell Technologies, #34815) at a density of 1.8×10^6^ cells/well (6,000 cells/microwell) in 1.5 mL/well of mTeSR Plus medium supplemented with 10 μM ROCK-inhibitor Y-27632, (ROCKi, StemCell Technologies, #72302). After 24 hours, the entire medium was replaced with mTeSR Plus Medium without ROCKi. From day 2 until day 21, the differentiation medium used was growth-factor-free chemically defined medium (gfCDM) (Muguruma et al. 2015), consisting of Isocove’s modified Dulbecco’s medium (Gibco, #12440053)/Ham’s F-12 (Gibco, #11765054) 1:1, chemically defined lipid concentrate (1% v/v, Gibco, #11905031), monothioglycerol (450μM, Sigma, #M6145), apo-transferrin (15 μg/ml, Sigma, #T1147), BSA (5 mg/ml, >99%, Sigma, #A9418), and 50 U/ml penicillin/50 μg/ml streptomycin (PS, Gibco, #15070063). The medium was also supplemented with insulin (7 μg/mL, Sigma, #91077C). At days 2 and 5, all medium was replaced for gfCDM supplemented with recombinant human basic FGF (FGF2, 50 ng/mL, Gibco, #13256029) and transforming growth factor β inhibitor (TGF-βi) SB431542 (SB, 10 μM, Sigma, #616461). On day 7, floating aggregates were transferred from microwell plates to ultra-low attachment 6-well plates and cultured at 1×10^6^ cells/ml in 1.8 mL/well, and 2/3 of the initial amount of FGF2 and SB was added. On day 14 and 18 entire medium was changed and FGF19 (100 ng/mL, PeproTech, #100-32) was added. From day 21, the aggregates were cultured in Neurobasal medium (Gibco, #12348017) supplemented with GlutaMax I (Gibco, #35050061s), N2 supplement (Gibco, #17502048), and PS. Recombinant human SDF1 (300 ng/mL, PreproTech, #300-28A) was added to culture on day 28.

### Maturation of cerebellar neurons

This part was performed as described in (Silva et al. 2020), with some modifications, namely the inclusion of N2B27 medium from days 35 to 64, instead of BrainPhys^TM^ Neuronal Medium as outlined in the original protocol. In short, day 35 aggregates were dissociated using Accutase and replated on wells coated with poly-L-ornithine (15 μg/mL, Sigma, #P3655) and Laminin (20 μg/mL, Sigma, #L2020), at a seeding density between 20,000 – 80,000 cells/cm^2^. Re-plated aggregates were cultured in N2B27 medium supplemented with Recombinant Human Brain Derived Neurotrophic Factor (BDNF, 20 ng/mL, PeproTech, #450-02) and Recombinant Human Glial-Derived Neurotrophic Factor (GDNF, 20 ng/mL, PeproTech, #450-10). N2B27 medium is a 1:1 mixture of DMEM:F12 (Gibco, #11320033) and Neurobasal medium supplemented with 0.5%v/v N2, 1%v/v B27 (Gibco, #17504044), 2 mM GlutaMax and 0.5%v/v PS. At day 64, BrainPhys^TM^ Neuronal Medium (StemCell Technologies, #05790) supplemented with NeuroCult^TM^ SM1 Neuronal Supplement (StemCell Technologies, #05711), N2 Supplement-A (StemCell Technologies, #07152), BDNF, GDNF, dibutyryl cAMP (1 mM, Sigma, #D0627), and ascorbic acid (200 nM, Sigma, #A92902) was added to the culture. Every 2-3 days, one-third of the total volume was changed.

### Organoid size analysis

To quantify organoid size over time in culture, several images were acquired at different time points using a DMI 3000 B (Leica) with a digital camera DXM 1200F (Nikon). The organoid diameter was calculated using Fiji-ImageJ, based on Feret’s diameter, the longest distance between any two points along the region of interest.

### Immunofluorescence

Different IF procedures were employed for organoids and cells plated on coverslips. For organoids, they were fixed at 4% (w/v) paraformaldehyde (PFA, Sigma, #P6148) for 45 min at 4°C in an agitation platform followed by two rinses in Phosphate buffered saline (PBS, 0.1M, Gibco, #21600-044) and overnight incubation in 15% (w/v) sucrose (Sigma, #S9378) at 4°C. Organoids were then embedded in 1:1 7.5% gelatin (Sigma, #G2500):15% sucrose (w/v) for 2h at 37°C and frozen in isopentane (Carlo Erba, #528492) at −80°C. Afterwards, organoids were cut in 12 μm sections on a cryostat-microtome (Leica CM3050S, Leica Microsystems), collected on Superfrost^TM^ Microscope Slides (VWR, #48311-703), and stored at −20°C. Sections were de-gelatinized for 45 min in PBS at 37°C. Then sections were incubated in 0.1 M Glycine (Sigma, #G8898) for 10 min at room temperature (RT), permeabilized with 0.1% v/v Triton X-100 (Sigma, #T9284) for 10 min at RT and blocked with 10% (v/v) fetal bovine serum (FBS, Gibco, #A5256701) in TBST [20 mM Tris-HCl pH 8.0, 150 mM NaCl, 0.05% (v/v) Tween-20)] for 30 min at RT. Primary antibodies were incubated 2h at RT or 4°C overnight (Table S3) diluted in blocking solution. Secondary antibodies were added to sections for 30 min (Table S4) at RT. 4’,6-diamidino-2-phenylindole (DAPI, 1.5 μg/mL, Sigma, #D9542) was used for nuclear counterstaining. Sections were mounted in Mowiol (Sigma, #10849).

In the case of plated cells on coverslips, they were washed twice with PBS 1x, fixed with 4% (w/v) PFA for 20 min at RT and kept at PBS until use. Then cells were incubated in 0.1 M Glycine for 10 min at RT, permeabilized with 0.1% (v/v) Triton X-100 for 10 min at RT and blocked with 10% (v/v) normal goat serum (NGS, Gibco, #PCN5000) in TBST (20 mM Tris-HCl pH 8.0, 150 mM NaCl, 0.05% Tween-20) for 30 min at RT. Primary and secondary antibodies were used the same way as for cryostat sections. For phalloidin staining, cells were incubated with Alexa Fluor 488 Phalloidin (1:40, Invitrogen, #A12379). Coverslips were mounted in Mowiol.

Microscopy experiments were performed at the Bioimaging Platform of the Gulbenkian Institute for Molecular Medicine. Fluorescent images were acquired using Zeiss LSM 710 confocal point-scanning microscope. Images were treated with ZEN 2012 SP5 FP3 (black; version 3.10) and Adobe Photoshop softwares.

### Quantification of immunostaining

Fluorescent images were quantified using ImageJ software. After the splitting of channels and automatic adjustment of a threshold, signal area was calculated. For each protein of interest, the ratio between its signal area and the nuclear DAPI staining, representing the total number of cells, was calculated, and converted to a percentage. The quantification of loop parameters in neural rosette structures was completed based on DAPI staining. To quantify the loop diameter three measurements for each ventricular structure were performed, pointing to the nearest pial surface, and the mean was calculated. The loop area was defined as the ratio between the loop area and the ventricle area. The apical and basal membrane lengths were also measured by ImageJ software. For dendritic spine classification, representative images with phalloidin staining were used to manually measure the number of spines, dendritic spine head width and neck length. Dendritic spines were classified as follows: Filopodia (length >2 μm); Long thin (length <2 μm); Thin (length <1 μm); Stubby (length/width ratio < 1μm); Mushroom (width >0.6μm) and Branched (2 or more heads) (Risher et al., 2014).

### COmbined Bisulfite Restriction Analysis (COBRA)

Genomic DNA from AS (AG1.0) and control (F002.1A.13) iPSCs at day 0 and day 80 of cerebellar differentiation were purified using conventional phenol:chloroform:isoamyl alcohol (Invitrogen, #15593-031) extraction, followed by bisulfite treatment using the EZ DNA Methylation Gold Kit (Zymo Research, #D5006). Bisulfite-treated DNA was amplified by a nested PCR approach for the PWS-IC region using the PCR primers (Table S5) with the KAPA HiFi Hot Start Uracil+ kit (Roche, #11828665001). PCR products were excised from the agarose gel using the NZYGelpure kit (NZYTech, MB01102) and then submitted to a restriction digestion with *BstU*I/*Bsh1236*I (Thermo Scientific, #FD0924) to distinguish methylated from unmethylated alleles based on the original status of the CGCG sequence using a previously validated assay (Pólvora-Brandao et al., 2018).

### Quantitative real time polymerase chain reaction (RT-qPCR)

Adherent cells and organoids were collected at different time points along the differentiation and dissociated with accutase during 7 min at 37°C. The enzyme was inactivated by addition of serum-containing medium and the cell pellet was resuspended in lysis/extraction buffer from the High Pure RNA Isolation Kit (Roche, 11828665001) and frozen at −80°C until further use. Total RNA was extracted following user’s manual instructions and converted into complementary cDNA with Transcriptor High Fidelity cDNA Synthesis Kit (Life Technologies, 4368814). SYBR Green Master Mix (Nyztech, MB44002) was used to analyze mRNAs of genes of interest (Table S5). All PCR reactions were done in duplicate or triplicate using the StepOneTM or ViiA™ 7 RT-PCR Systems (Applied BioSystems). Results were analysed with StepOne^TM^ or QuantStudio™ RT-PCR Software. Quantification was performed by calculating the ΔCt value using glyceraldehyde-3-phosphate dehydrogenase (*GAPDH)* as housekeeping gene and results are presented as mRNA expression levels (2^−ΔCt^) relative to *GAPDH*.

### Pyrosequencing

Allelic expression analysis was carried out on an iPSC line derived from a patient carrying a defined three-base pair deletion in maternal *UBE3A* allele (AS_Δ3) (Stanurova et al. 2016). Total RNA was extracted at different time points of the differentiation process (days 0, 35, 56 and 80) and retrotranscribed into cDNA as stated in the RT-qPCR section. For pyrosequencing, PCR and sequencing primers were designed using the PyroMark Assay Design 2.0.1.15 software (Qiagen, score 100) (Table S5). PCR amplification of maternal and paternal *UBE3A* alleles was analysed by pyrosequencing with the PyroMark Q24 workstation according to the manufacturer’s instructions (Qiagen) and the relative level of the two alleles was quantified by the PyroMark Q24 2.0.07 Software. Primer sequences are in Table S5.

### Flow cytometry

Aggregates were dissociated using Accutase and fixed drop-by-drop with 70%v/v Ethanol (chilled at −20°C, Fisher, #E/0650DF/C17) with vortexing and stored at −20°C. Eppendorf tubes were coated with 1%w/v BSA in PBS for 15 min, before adding samples, centrifuging at 1000 rpm for 10 minutes, and washing cells twice with PBS. For intracellular staining, cells were resuspended in 3%w/v BSA in PBS, permeabilized with 0.1%w/v saponin (Sigma, #47036) in 3%v/v FBS for 15 minutes at RT. Cells were washed twice with 1%v/v FBS and incubated with primary mouse anti-Ki67 antibody (1:20; BD Biosciences, #571599 RRID:AB_3686606) diluted in 3% FBS for 1 hour at RT. Cells were washed three times with 1%v/v FBS and incubated with secondary goat anti-mouse AlexaFluor-488 antibody (1:1000; Invitrogen, #A-11001 RRID:AB_2534069) in 3%v/v for 45 minutes at RT in the dark. Cells were washed and resuspended in PBS. For the apoptosis / necrosis assay (Abcam, #ab176750), live cells from dissociated aggregates were resuspended in Assay Buffer and stained with Apopxin Deep Red, Nuclear Green DCS1 and Cytocalcein Violet 450 for 30 minutes in the dark, following manufacturer’s recommendations.

Samples were analysed in a FACSCalibur™ flow cytometer (Becton Dickinson), and for each sample, 10,000 events were recorded within the defined gate based on forward (FSC) and side scatter (SSC). Gates were selected to contain less than 1% of false positives, and all results were analysed using the FlowJo software.

### Single cell calcium imaging

For single cell calcium imaging, organoids were dissociated using accutase and re-plated on glass bottom microwell chambers (MatTek, #CCS-8) previously coated with poly-D-lysine (Sigma, #P1024) and Laminin (20 μg/mL), at a seeding density of 80,000 cells/cm^2^. At different time-points of differentiation, cells were loaded with 5 μM Fura-2 AM (Invitrogen, #F1201) in Krebs solution (132 mM NaCl, 4 mM KCl, 1.4 mM MgCl2, 2.5 mM CaCl2, 6 mM glucose, 10mM HEPES, pH 7.4) for 45 min at 37°C. Microwell chambers were washed in Krebs solution and then mounted on an inverted microscope with epifluorescence optics (Axiovert 135TV, Zeiss). Cells were continuously perfused with Krebs solution and stimulated by applying high-potassium Krebs solution (KCl, 50 mM) and Histamine (100 μM), as previously described (Agasse et al. 2008; Xapelli et al. 2013). The time course of experiments was the following: 5 min baseline, stimulation with KCl from 5 to 7 min, KCl washout from 7 min to 10 min, stimulation with histamine from 10 to 12 min and histamine washout from 12 to 15 min. Ratio images were obtained from image pairs acquired every 200 ms by exciting the cells at 340 nm and 380 nm. Excitation wavelengths were changed through a high-speed switcher (Lambda DG4, Sutter Instrument, Novato, CA, United States). The emission fluorescence was recorded at 510 nm by a cooled CDD camera (Photometrics CoolSNAP fx). Images were processed and analysed using the software MetaFluor (version 1.0.93, Universal Imaging, West Chester, PA, United States). Regions of interest were defined manually over the cell profile. KCl and histamine peaks given by the normalized ratios of fluorescence at 340/380 nm, at the proper time periods, were used to calculate the ratios of the responses.

### Patch-Clamp Recordings

Whole-cell patch-clamp recordings were obtained from cerebellar neurons visualized with an upright microscope (Zeiss Axioskop 2FS) equipped with differential interference contrast optics using a Zeiss AxioCam MRm camera and a x40 IR-Achroplan objective. During recordings, cells were continuously superfused with artificial cerebrospinal fluid (aCSF) containing 124 mM NaCl, 3 mM KCl, 1.2 mM NaH2PO4, 25 mM NaHCO3, 2 mM CaCl2, 1 mM MgSO4, and 10 mM glucose, which was continuously gassed with 95% O2/5% CO2. Recordings were performed at RT in current-clamp or voltage-clamp mode [holding potential (Vh) = −70 mV] with an Axopatch 200B (Axon Instruments) amplifier, as performed elsewhere (Felix-Oliveira et al. 2014). Synaptic currents and action potential activity were recorded using patch pipettes with 4–7 MΩ resistance filled with an internal solution containing 125 mM K-gluconate, 11 mM KCl, 0.1 mM CaCl2, 2 mM MgCl2, 1 mM EGTA, 10 mM HEPES, 2 mM MgATP, 0.3 mM NaGTP, and 10 mM phosphocreatine, pH 7.3, adjusted with 1 M NaOH, 280-290 mOsm. Acquired signals were filtered using an in-built, 2-kHz, three-pole Bessel filter, and data were digitized at 5 kHz under control of the pCLAMP 11 software program. The junction potential was not compensated for, and offset potentials were nulled before gigaseal formation. The resting membrane potential was measured immediately upon establishing whole-cell configuration and the junction potential of the electrode was considered (−12 mV).

For each neuron, the threshold for action potential generation was determined as the difference between the resting membrane potential and the membrane potential at which phase plot slope reached 10 mV/ms (Naundorf et al., 2005). Analysis of the amplitude and half-peak within were performed offline using the Clampfit 11 software. In the voltage-clamp mode, spontaneous miniature postsynaptic currents were recorded in a CSF solution during 5 min. Analysis of miniature currents was evaluated using the Mini-Analysis Software.

### Bulk RNA-sequencing

Total RNA from iPSC-derived day 56 neurons of control (F002.1A.13, n=2; iPSC6.2, n=1) and AS (AG 1.0 n=2, ASD n=1) was extracted using the High Pure RNA Isolation Kit. Total RNA was then depleted of ribosomal RNA, and strand-specific libraries were prepared and sequenced using Illumina paired-end protocols with a 150 bp sequencing length (Novogene Co., LTD). Raw reads were quality controlled using FastQC (Babraham Institute n.d.; v.0.11.5), and sequencing adapters removed using trim_galore (Babraham Institute n.d.; v.0.6.6). Paired-end alignments were generated using STAR (Dobin et al., 2012) (v.2.7.5a) and the human reference genome (hg38/GRCh38), using parameters that allow the assignment of multiple-mapped reads to a random position (--outFilterMultimapNmax 5000, -- outSAMmultNmax 1, --outFilterMismatchNmax 999, --outFilterMismatchNoverLmax 0.06). Reads covering exons of each annotated gene were counted using featureCounts (Liao et al., 2013) (v.1.6.0) and the comprehensive gene annotation from Gencode (Frankish et al., 2019) v40 (GRCh38.13), using parameters suitable to count multi-mapped reads (-M -p -C -s 2).

### Normalization and Differential Expression Analysis

Normalization and differential expression analysis were performed using the R package DESeq2 (Love et al., 2014) (v.1.32.0). DESeq2 applies its internal median-of-rations method to normalise for sequencing depth and RNA composition bias, especially in cases when only a small number of genes are very highly expressed in one condition but not another.

For differential expression analysis, DESeq2 uses a Wald test for hypothesis testing to compare multiple groups and test significance. *P*-values were corrected for multiple testing using the Benjamini-Hochberg false-discovery-rate (FDR) procedure. For principal components analysis (PCA, computed using the PCAtools (Blighe, 2021) library), raw counts were normalized using the variance stabilizing transformation (VST) from DESeq2.

Gene ontology (GO) enrichment analysis was performed using the gseGO() function from the clusterProfiler R package (v4.2.2), applying Gene Set Enrichment Analysis (GSEA) to genes ranked by their Wald test statistics from DESeq2 differential expression results. Enrichment was tested across all three GO domains (Biological Process, Molecular Function, Cellular Component) with the following parameters: minGSSize = 3, maxGSSize = 800, pvalueCutoff = 0.05, and Benjamini–Hochberg FDR correction (pAdjustMethod = “BH”).

Genes ranked according to the value for the Wald statistic were analysed for Gene Set Enrichment Analysis (Mootha et al., 2003; Subramanian et al., 2005) using the R package fgsea (Korotkevich et al., 2021). For GSEA, we used gene sets retrieved from the MSigDB (Liberzon et al., 2015; Liberzon et al., 2011). Associations with False Discovery Rate (FDR) lower than 5% were considered significant.

All the analysis were performed in RStudio (v1.4.1717), using R (v4.1.2). Besides the Bioconductor packages for RNA-seq analysis, the following packages were used: pheatmap (v.1.0.12), reshape2 (v1.4.4), ggplot2 (v.3.3.6), RColorBrewer (v.1.1-3), magrittr (v.2.0.3), vsn (v.3.62.0), ggrepel (v.0.9.1), data.table (v.1.14.2), tidyverse (v.1.3.1), eulerr (v.6.1.1), UpSetR (v.1.4.0), ggpubr (v.0.4.0), PCAtools (v.2.6.0), clusterProfiler (v.4.2.2), enrichplot (v.1.14.2) and fgsea (v.1.20.0).

For GO terms analysis, genes were considered as significantly differentially expressed when the adjusted p-values were < 0.01, and the log2 of the fold change was < ( −1) for downregulated genes or > 1 for upregulated genes. For all the other analyses, genes were considered as significantly differentially expressed when the adjusted p-values were < 0.05, and the log2 of the fold change was < ( −1) for downregulated genes or > 1 for upregulated genes.

### Statistical Analysis

Quantitative data were generated at least in biological triplicates per experimental group. Data were analysed by using GraphPad Prism 8.4.3 (GraphPad Software, San Diego, CA, USA) software. All results are presented as the mean ± SEM of the number of experiments performed in replicates as indicated in the figure legends. Normal distribution was assessed by the Kolmogorov-Smirnov test; comparisons between two groups were performed by unpaired two-tailed Student’s *t*-test or non-parametric Mann-Whitney test; comparisons among multiple groups were performed by two-way ANOVA followed by Sidak’s or Tukey’s multiple comparisons tests. Significance was defined as p<0.05; Statistically significant differences are documented in the respective figure legend.

