## Supplemental Figures for "Stem-cell modeling of cerebellar dysfunction of Angelman syndrome"

### SUPPLEMENTARY FIGURES

Figure S1

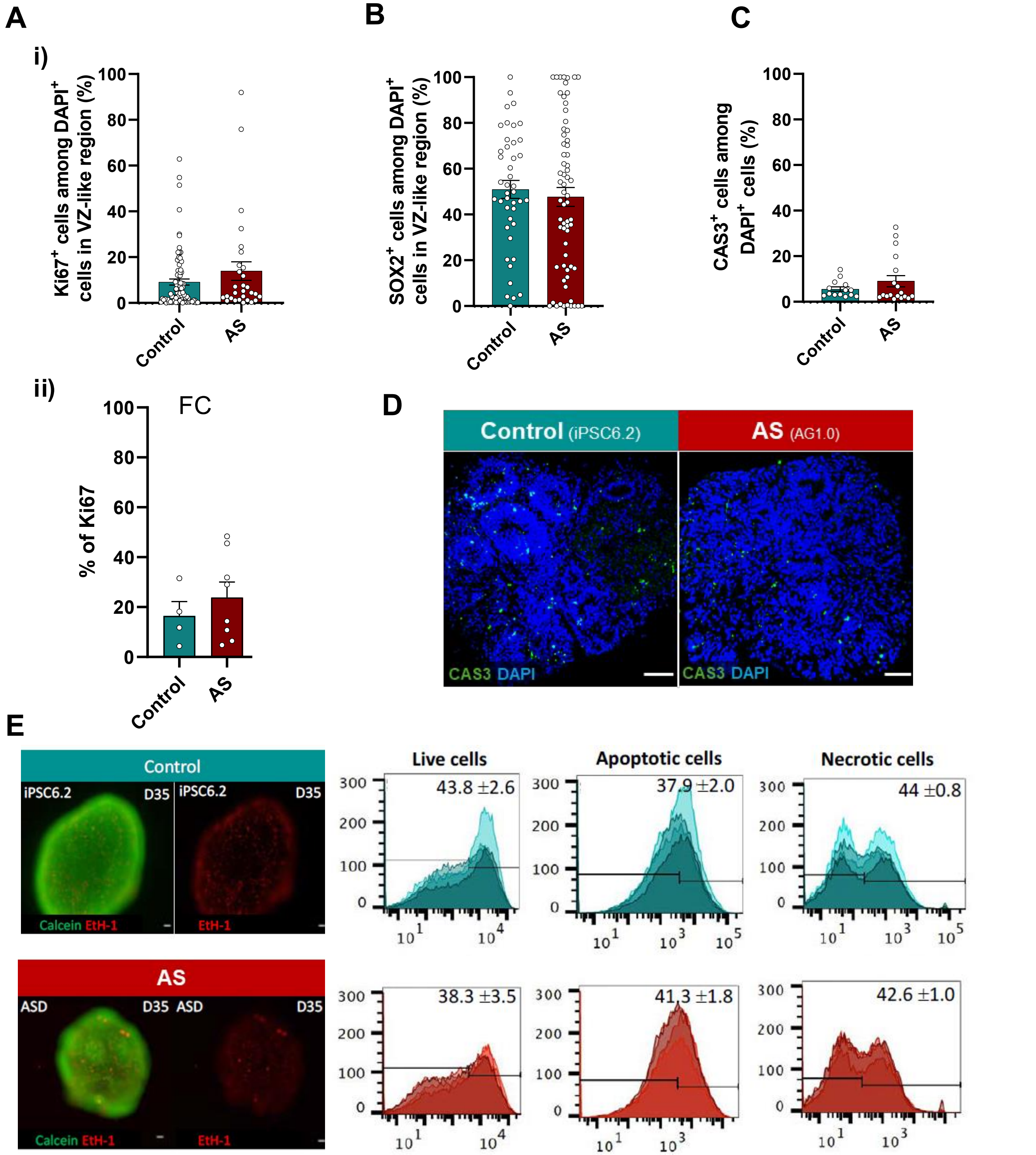
A
i)
Ki67+ cells among DAPI+ cells in VZ-like region (%)
0
20
40
60
80
100
Control
AS
B
SOX2+ cells among DAPI+ cells in VZ-like region (%)
C
CAS3+ cells among DAPI+ cells (%)
ii)
FC
% of Ki67
D
Control (iPSC6.2)
AS (AG1.0)
CAS3 DAPI
E
Control
iPSC6.2
D35
Calcein EtH-1
EtH-1
AS
ASD
Live cells
Apoptotic cells
Necrotic cells
43.8 ±2.6
37.9 ±2.0
44 ±0.8
38.3 ±3.5
41.3 ±1.8
42.6 ±1.0
300
200
100
0

**Figure S2**

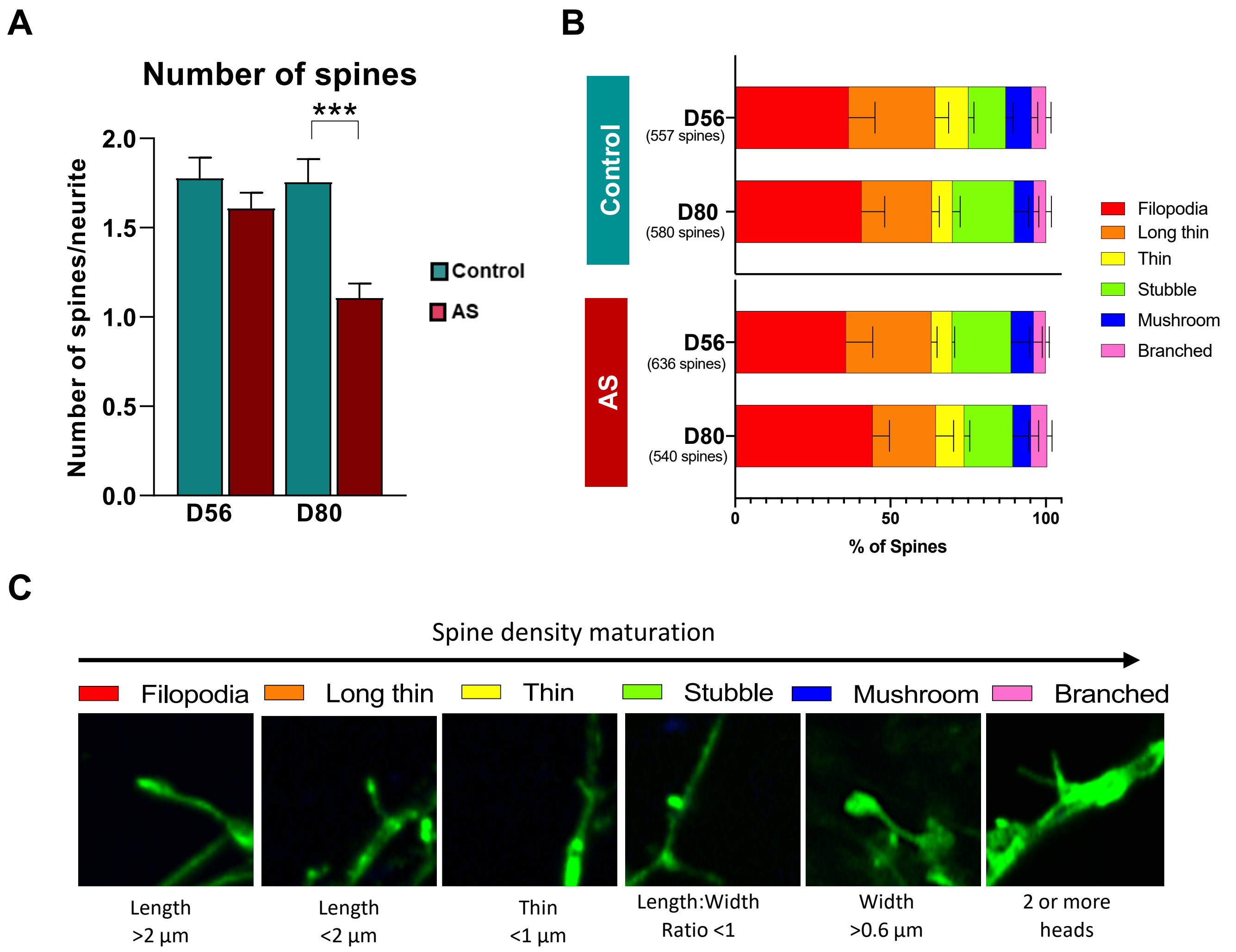
A
Number of spines
***
2.0
1.5
1.0
0.5
0.0
Number of spines/neurite
D56
D80
Control
AS
B
Control
AS
D56
(557 spines)
D80
(580 spines)
D56
(636 spines)
D80
(540 spines)
0
50
100
% of Spines
Filopodia
Long thin
Thin
Stubble
Mushroom
Branched
C
Spine density maturation
Filopodia
Long thin
Thin
Stubble
Mushroom
Branched
Length >2 µm
Length <2 µm
Thin <1 µm
Length:Width Ratio <1
Width >0.6 µm
2 or more heads

**Figure S3**

**A**

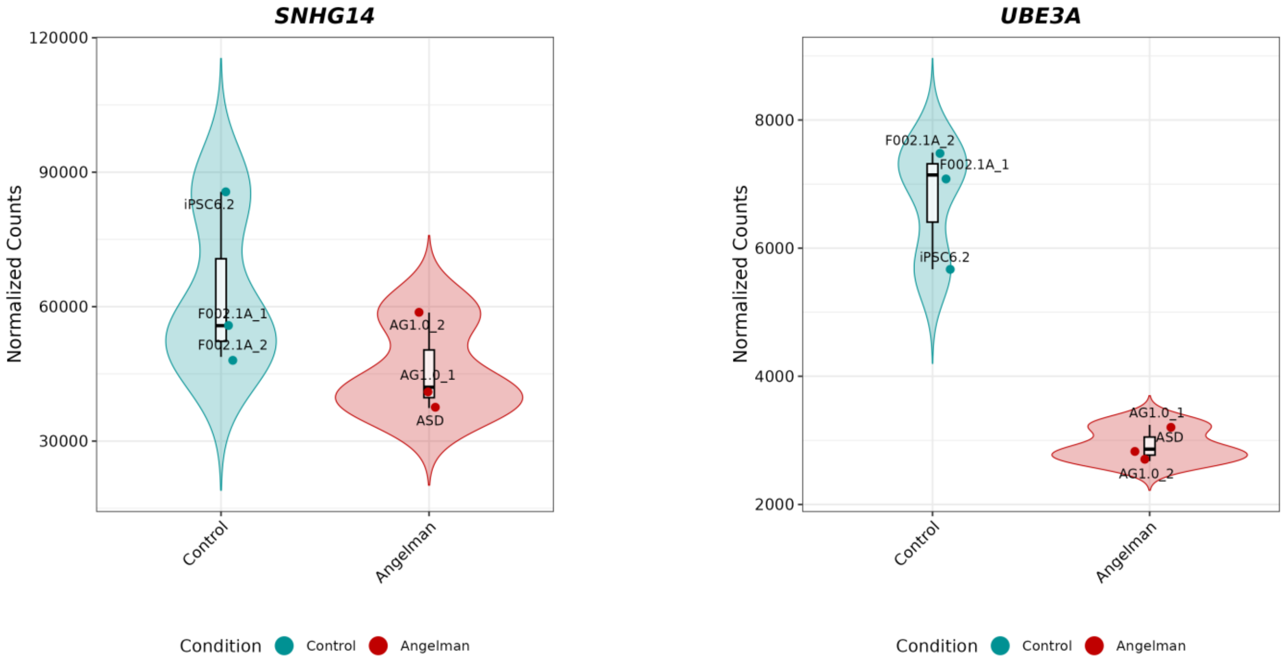

SNHG14
UBE3A
Normalized Counts
120000
90000
60000
30000
8000
6000
4000
2000
iPSC6.2
F002.1A_1
F002.1A_2
AG1.0_2
AG1.0_1
ASD
Control
Angelman
Condition


**B**

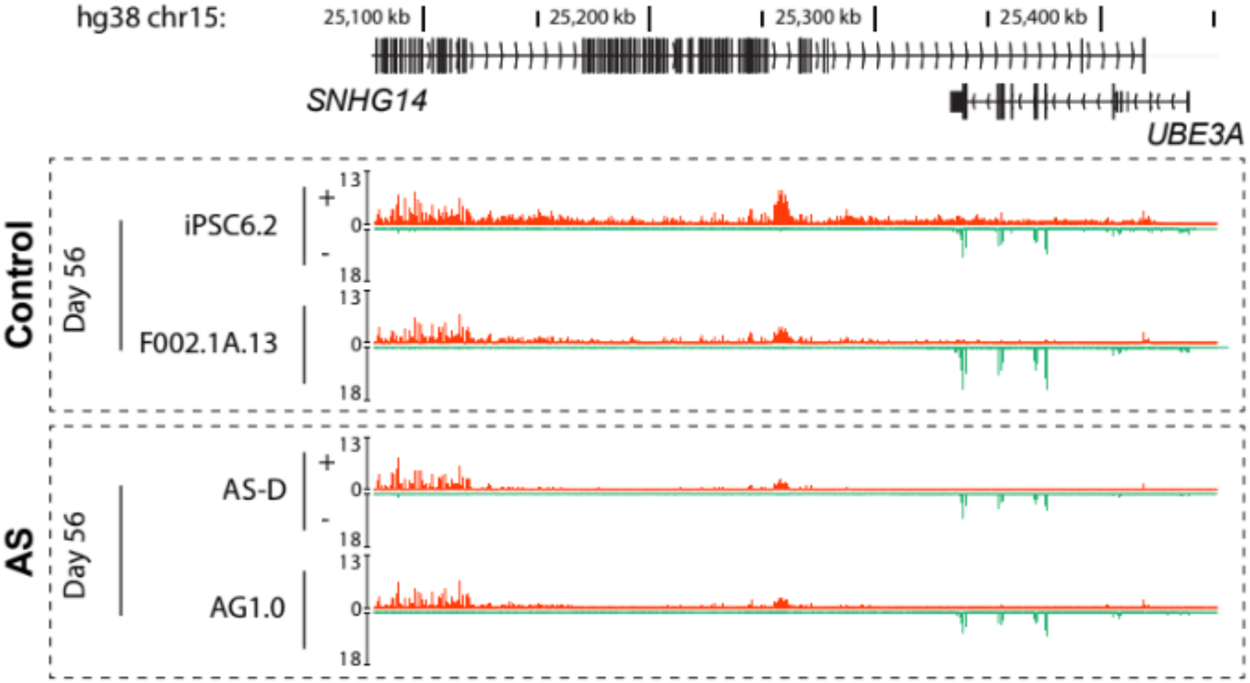

hg38 chr15:
25,100 kb
25,200 kb
25,300 kb
25,400 kb
SNHG14
UBE3A
Control
Day 56
iPSC6.2
F002.1A.13
AS
Day 56
AS-D
AG1.0
13
0
18
+
-

**Supplementary Figure 1. Proliferation and apoptosis in control and AS-derived cerebellar organoids.**

**A)** ***i)*** Quantification of Ki67 positive nuclei in ventricular zone (VZ)-like regions of cerebellar organoids on day 35 in AS (n=29) vs. unaffected control (n=86) day 35 cerebellar organoids. ***ii)*** Flow cytometry (FC) analysis of Ki67 in AS (n=8) vs. unaffected control (n=4) day 35 cerebellar organoids. Data from both AS (AG1.0 & ASD) and unaffected control (F002.1A.13 & iPSC6.2) lines were combined.

**B)** Quantification of the percentage of SOX2 positive cells in the ventricular zone (VZ)-like region in AS (n=66) vs. unaffected control (n=43) day 35 cerebellar organoids. Data from both AS (AG1.0 & ASD) and unaffected control (F002.1A.13 & iPSC6.2) lines were combined.

**C)** Quantification of CAS3-positive cells as a proportion of total DAPI-stained nuclei in AS (AG1.0; n=18) vs. unaffected control (iPSC6.2; n= 14) day 35 cerebellar organoids.

**D)** Representative images of immunostaining for phosphorylated caspase 3 (CAS3; in green) in AS (AG1.0) and unaffected control (iPSC6.2) cerebellar neurons. Nuclei counterstained with DAPI. Scale bars, 50 µm.

**E)** Fluorescence viability analysis. On the left, calcein_EtH-1 staining of live organoids [calcein AM for live cells in green; ethidium homodimer-1 (EtH-1) for dead cells in red]. On the right, histograms of live (Cytocalcein Violet 450), apoptotic (Apopxin Deep Red) and necrotic (Nuclear Green DCS1) cells evaluated by flow cytometry analysis. Data are represented as mean ± SEM of n = 4 independent experiments, data from the two independent AS (AG1.0 & ASD) and control (iPSC6.2 & F002.1A.13) iPSC lines were pooled for analysis.

**Supplementary Figure 2. Spine morphology in control and AS-derived neurons at day 56 and 80.**

**A)** Quantification of the number of spines per neurite in control and AS-derived neurons at days 56 (AS n=407, Ctrl n=277) and 80 (AS n=272, Ctrl n=301), data from the two independent AS (AG1.0 & ASD) and control (iPSC6.2 & F002.1A.13) iPSC lines were pooled for analysis. Spines were stained with phalloidin. Statistically significance represented as ***p<0.001, two-way ANOVA test followed by Sidak's multiple comparisons test. two-tailed Student's t-test (two-tailed) statistics: ***p<0.001.

**B)** Quantification of dendritic spine morphologies observed on days 56 and 80. Total number of spines analysed is indicated on the graph. Control: n=4; AS: n=5/6; data from the two independent AS (AG1.0 & ASD) and control (iPSC6.2 & F002.1A.13) iPSC lines were pooled for analysis.

**C)** Representative phalloidin-stained images illustrating the range of dendritic spine morphologies analysed.

**Supplementary Figure 3 - Reduced *UBE3A* expression in AS cerebellar neurons**

**A)** Violin plots showing normalised RNA-seq counts for *SNHG14* and *UBE3A* in day 56 cerebellar cultures derived from unaffected control (iPSC6.2, n=1; F002.1A.13, n=2) and AS (ASD, n=1; AG1.0, n=2) cerebellar neurons at day 56 of differentiation.

**B)** Genome browser views of the *SNHG14* and *UBE3A* locus on chromosome 15 (hg38), illustrating strand-specific RNA-seq coverage in control (iPSC6.2 & F002.1A.13) and AS (ASD & AG1.0) cerebellar neurons at day 56 of differentiation. Orange tracks indicate reads mapped to the forward strand (primarily *SNHG14*), while green tracks correspond to reads on the reverse strand, belonging to the *UBE3A* transcript.
